# Optimally designed vs intuition-driven inputs: the study case of promoter activity modelling

**DOI:** 10.1101/346379

**Authors:** L. Bandiera, V. Kothamachu, E. Balsa-Canto, P. S. Swain, F. Menolascina

**Affiliations:** School of Engineering, Institute for Bioengineering, The University of Edinburgh, Edinburgh, E119 3DW, UK.; SynthSys - Centre for Synthetic and Systems Biology, The University of Edinburgh, Edinburgh, E119 3BF.; (Bio)Process Engineering Group, IIM-CSIC Spanish National Research Council, Vigo, Spain.

## Abstract

Synthetic biology is an emerging engineering discipline that aims at synthesising logical circuits into cells to accomplish new functions. Despite a thriving community and some notable successes, the basic task of assembling predictable gene circuits is still a key challenge. Mathematical models are uniquely suited to help solve this issue. Yet in biology they are perceived as expensive and laborious to obtain because *low-information* experiments have often been used to infer model parameters. How much additional information can be gained using optimally designed experiments? To tackle this question we consider a building block in Synthetic Biology, an inducible promoter in yeast *S. cerevisiae*. Using *in vivo* data we re-fit a mathematical model for such a system; we then compare *in silico* the quality of the parameter estimates when model calibration is done using typical (e.g. step inputs) and optimally designed experiments. We find that Optimal Experimental Design leads to ~70% improvement in the predictive ability of the inferred models. We conclude providing suggestions on how optimally designed experiments can be implemented *in vivo*.

## I. INTRODUCTION

Synthetic Biology is an emerging discipline that seeks to implement *de novo* tasks in cells. Despite a booming community and the opportunities that Synthetic Biology offers [1], the assembly of synthetic circuits with predictable functions remains a challenge. If Synthetic Biology is to advance towards application, it is necessary to increase the predictability of gene network dynamics. Mathematical models offer a means to achieve this goal, yet their use in Synthetic Biology has so far only been limited [2].

The reason for the low adoption of models largely lies in the limitations of traditional experimental platforms in biology (e.g. microplate readers). “Pulses” or “steps” of chemicals are the *de facto* standard stimuli to probe cell behaviour. Such designs, however, often allow for poorly-informative experiments. Indeed, chemical “steps” and “pulses” are low-pass filtered by molecular diffusion, which limits their frequency content. Furthermore, the inherent non-linearity of biological networks prevents the adoption of results on *persistent excitation* developed for the identification of linear models [3].

These facts raise the questions of whether it is possible to design informative experiments for the identification of gene networks and how to do so.

Model-based Optimal Experimental Design (MBOED) allows the design of maximally informative experiments and has recently been adopted in the identification of biological systems. For example, Bandara *et al.* showed that optimally designed (yet technologically constrained) experiments lead to a 60-fold reduction in the mean variance of parameter estimates over experience-based schemes [4]. Similarly, Ruess *et al.* emphasised the improvement of optimised dynamic inputs over random stimulation patterns in the characterisation of a light-inducible promoter [5].

The adoption of MBOED in (Synthetic) Biology generally faces a large amount of inertia: optimally designed experiments are difficult to implement with traditional experimental platforms and the skills to design them are not widespread in wet laboratories. Technological developments (e.g. microfluidics) and computational tools (e.g. AMTGO2 [6]) allow this limitation to be overcome but they have steep learning curves. The question is then: does the gain in information OED offers justify the efforts of adopting it?

To address this question, here we consider the identification of a mathematical model of a building block in Synthetic Biology: an inducible promoter. Many synthetic promoters are available, but since they generally use DNA sequences from the same organisms they are engineered for, they suffer from unwanted regulation from other genes in the genome. This makes disentangling and modelling promoter activity a non-trivial task. To overcome this issue, we focus on an orthogonal promoter [7], i.e. a promoter built in a species (*S. cerevisiae*) using DNA sequences from a different one (*E. coli*). This promoter, designed by Gnügge *et al.* [7] (Fig. 1), drives the expression of a fluorescent reporter, Citrine, when cells are exposed to the chemical IPTG. IPTG enters the cell through the permease Lac12 and binds the LacI protein, thereby relieving its repression on the promoter activity. Binding of the constitutively expressed tTA to the tetO_2_ site results in expression of Citrine.

**Fig. 1:**
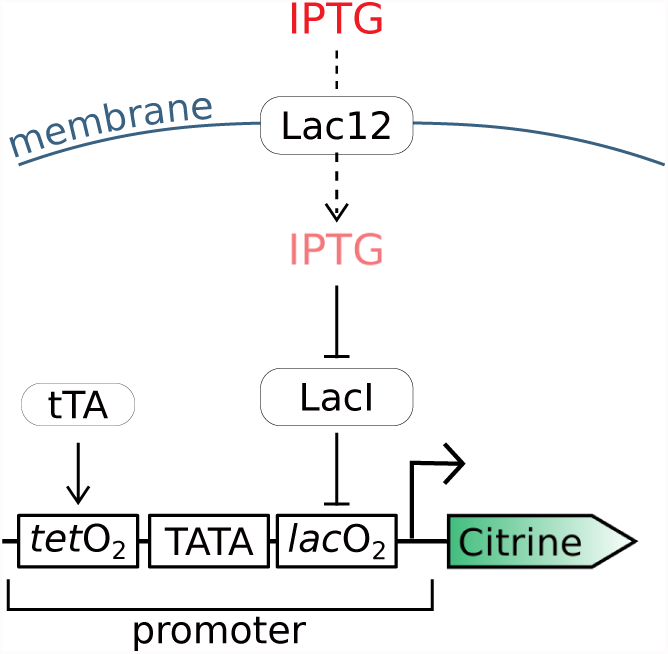
Schematic of the inducible promoter implemented in [7]. A native S. cerevisiae promoter was engineered by cloning the (tetO)_2_ and (lacO)_2_ operator sequences upstream and downstream of the TATA box respectively. The construct was integrated into the genome of a budding yeast strain constitutively expressing the heterologous transcription factors - tetracycline responsive transactivator (tTA) and LacI repressor, and the lactose permease (Lac12). The activity of the resulting promoter, regulated by the exogenous, non-metabolizable inducer β − D − 1 thiogalactopyranoside (IPTG), is reported by the expression of the Citrine fluorescent reporter.

Based on published characterisation data [7], we first refine a mathematical model of the inducible promoter (*M_PLac_*), obtaining *M_PLac_*_,*r*._ We then define a reduced model structure, 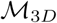, able to mimic the dynamics of *MP_Lac,r_* (*M_IP,r_*). We hence simulate the response of *M_IP,r_* to both optimal and intuition-driven inputs and compare the amount of information provided by each input class using the posterior distributions of the inferred parameters.

Not only do our results suggest that MBOED allows the design of more informative experiments for the characterisation of synthetic promoters, they also provide a conservative estimate of the improvement in parameter accuracy that can be achieved via MBOED.

The manuscript is organised as follows: Section II discusses how we recalibrate a model of the inducible promoter, define a lower-order model and use it to compare the informativeness of different input classes. Section III elaborates on the importance of OED for the design of more informative experiments in Synthetic Biology. Section IV details our *in silico* experiments, the comparison of informativeness of different input classes and the design of optimal experiments. Finally, Section V presents our conclusions and future directions.

## II. RESULTS

### A. Refitting Gnügge et al.’s Model

As starting point of our analysis we consider *M_PLac_*, the model proposed by Gnügge and colleagues [7]. We first seek to independently assess the ability of this model to capture the experimental data reported in the original paper [7], comprising several IPTG dose-response curves sampled at five equidistant time points after induction. We note that at intermediate concentrations model predictions appear to systematically underestimate the measured steady states by 20-30% (Fig. 2, grey line).

**Fig. 2:**
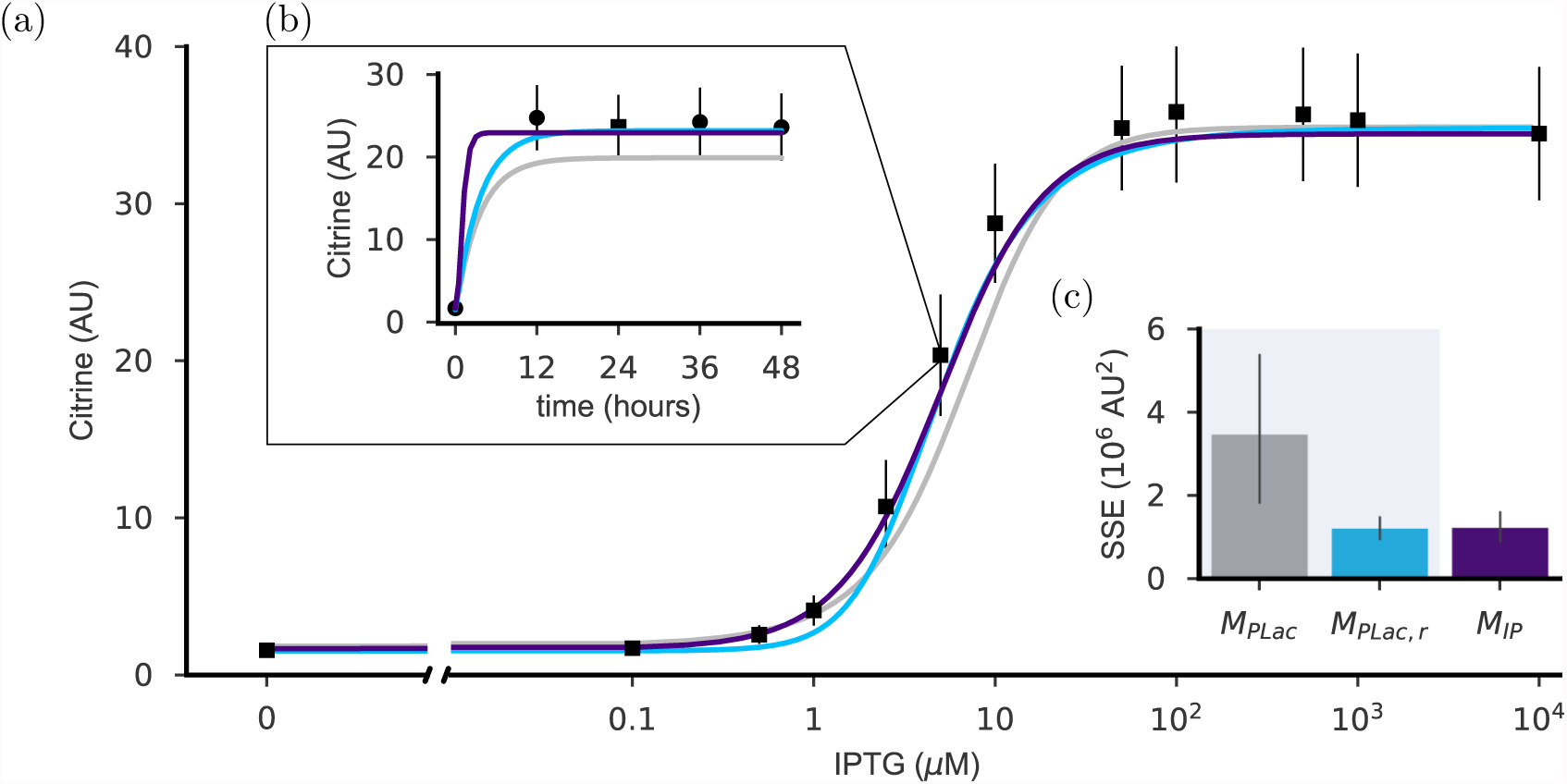
Comparison between *M_PLac_*, *M_PLac,r_* and *M_IP_* model structures. (a) Dose-response curve after 24 hours of incubation with the specified IPTG concentrations. Experimental data, median (filled squares) and inter-quartile range (errorbars) of Citrine distributions, were retrieved from [7]. Solid lines show the in-silico dose-response curve for M_PLac_ (grey), MP_Lac,r_ (cyan) and M_IP_ (purple). (b) The full data includes the dynamics of Citrine. An example is shown for the induction with 5 µM of IPTG. (c) Barplot of the sum of squared errors of predictions (SSE), quantifying the predicted deviations from empirical data.

Reasoning that this discrepancy would offer an opportunity to refine *M_PLac_*, we re-calibrate the model using enhanced Scatter Search (eSS) and obtain a new model, *M_PLac,r_* (Fig. 2a, cyan), that generally better fits the available experimental data (Fig. 2b). *M_PLac,r_* offers a 56% improvement in fit, as quantified by the sum of squared errors of predictions (SSE) (Fig. 2c).

### B. A reduced-order model captures the dynamics of the inducible promoter

To constrain the number of parameters to be identified and the computational cost associated to optimal experimental design, we develop a lower-order model structure (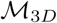). The model structure reads as follows:

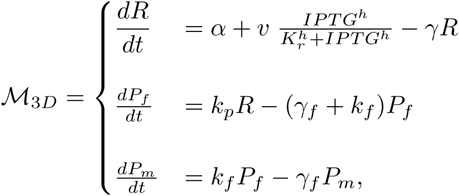

where *R*, *P_f_* and *P_m_* are the concentrations of Citrine mRNA, immature folded protein and matured (fluorescent) protein, respectively. The model features 8 parameters: *α* and *v* are the basal and maximal transcriptional rate respectively; *h*, the Hill coefficient; *K_r_*, the Michaelis-Menten coefficient; *k_p_*, the translation rate and the rate of maturation of the folded protein, *k_f_*. All biochemical species are subject to linear degradation, occurring at rates *γ* for mRNA and *γ_f_* for protein. This model structure builds on the assumption that the expression of LacI and Lac12, as well as the binding of LacI-dimer to the operator sites and to IPTG, occurs on faster time scales than Citrine expression. Fitting all parameters of 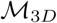 to the time-series data in [7], we obtain *M_I P_*.

When compared with *M_PLac_* and *M_PLac,r_,* we find that MIP best fits the measured steady-states, i.e. the dose-response curve (Fig. 2a), as well as the experimental data acquired at time-points different from 24 hours (see Fig. 2b for an example). Despite its lower order, MIP achieves predictive capabilities comparable to *M_PLac,r_*. To show this, we calculate the SSE over the whole set of experimental data (Fig. 2c). It is interesting to note that MIP is characterised by a smaller rise time (1.8 hours) than both *M_PLac_* and *M_PLac,r_* (7.9 hours) (Fig. 2b). Here, the rise time is defined as the time required for the output to rise from 10% to 90% of the steady-state. We also note that the long sampling intervals used in the original study [7] does not allow further constraining the characteristic time-scale of the system.

As we aim to compare the informative content of different input classes (II, section C), we need our reduced model to mimic as closely as possible the dynamics of the genetic system of interest. We therefore consider *M_PLac,r_* as our nominal model and generate a set of pseudo-experimental data to re-calibrate *M_IP_*; in so doing we obtain *M_IP,r_*. It is worth noting that we could use *M_IP_* for MBOED, however the ability to generate additional datasets and further constrain parameter calibration made us prefer *M_Lac,r_* as our reference. Considering the limited complexity of the underlying biological system, we decided to use stepwise, pulses, ramp wise and stepwise random inputs in the pseudo-data generation. The results of *M_IP_,_r_* identification show that this model recapitulates the dynamics of *M_PLac,r_* (Fig. 3).

**Fig. 3:**
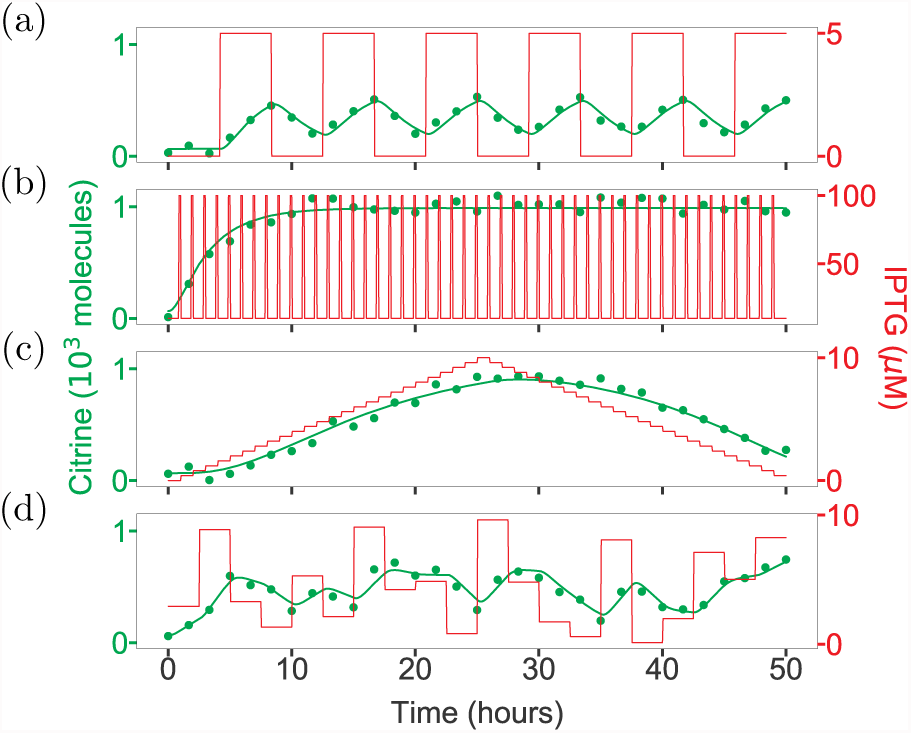
Pseudo-experiments for the identification of M_IP,r_. Step (a), pulse (b), ramp (c) and random (d) inputs (red line) were applied to M_PLac,r_ to simulate Citrine dynamics and to obtain pseudo-data (green circles). The response of the calibrated M_IP,r_ is shown as a green, solid line.

### C. Intuition-driven inputs are equally (poorly) informative

We first seek to compare the informative yield of intuition-driven stimuli (step, pulse and random) for the calibration of 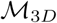. With this aim, we generated *N_j_* = 100 input profiles for each of the three classes (Methods, section IV-B). By simulating the output of *M_IP,r_* for each input, we obtain pseudo-experimental data we use for the calibration of 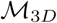. We formulate parameter estimation as a non-linear optimisation problem and use eSS to solve it. As the posterior distributions of parameter estimates are not Gaussian, we cannot use standard metrics (e.g. z-score) to assess the statistical significance of the distance between nominal and estimated parameter value. To overcome this limitation, we compute the *relative error* 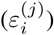 between each parameter estimate 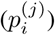 and its nominal value 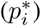:

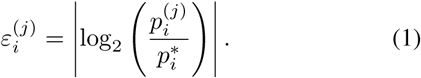

where *i* identifies the *i^th^* entry in the parameter vector and *j* is the index of the input profile yielding the parameter estimate 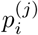. Notably, 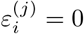 when the parameter estimate equals its nominal value, while the absolute value ensures that under and over estimates are treated equally.

The distributions of relative error (*ε_i_*) for the 100 input profiles highlight a differential sensitivity of the output to the parameters (Fig. 4b-d). It is worth noting that the high variability in the estimates of *α*, *v* and *γ* agrees with a preliminary identifiability analysis (results not shown), suggesting high correlation between these parameters. Practical identifiability issues have indeed the potential to hinder our ability to identify the affected parameters with high confidence. Overall, the *ε_i_* distributions suggest that the intuition-driven inputs convey a similar amount of information (Fig. 4b-d). This is further confirmed by the absence of a statistically significant difference in the *average relative error* 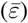 metric (Fig. 4f), defined as:

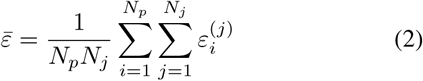

where *N_p_* is the number of parameters in the model structure.

**Fig. 4:**
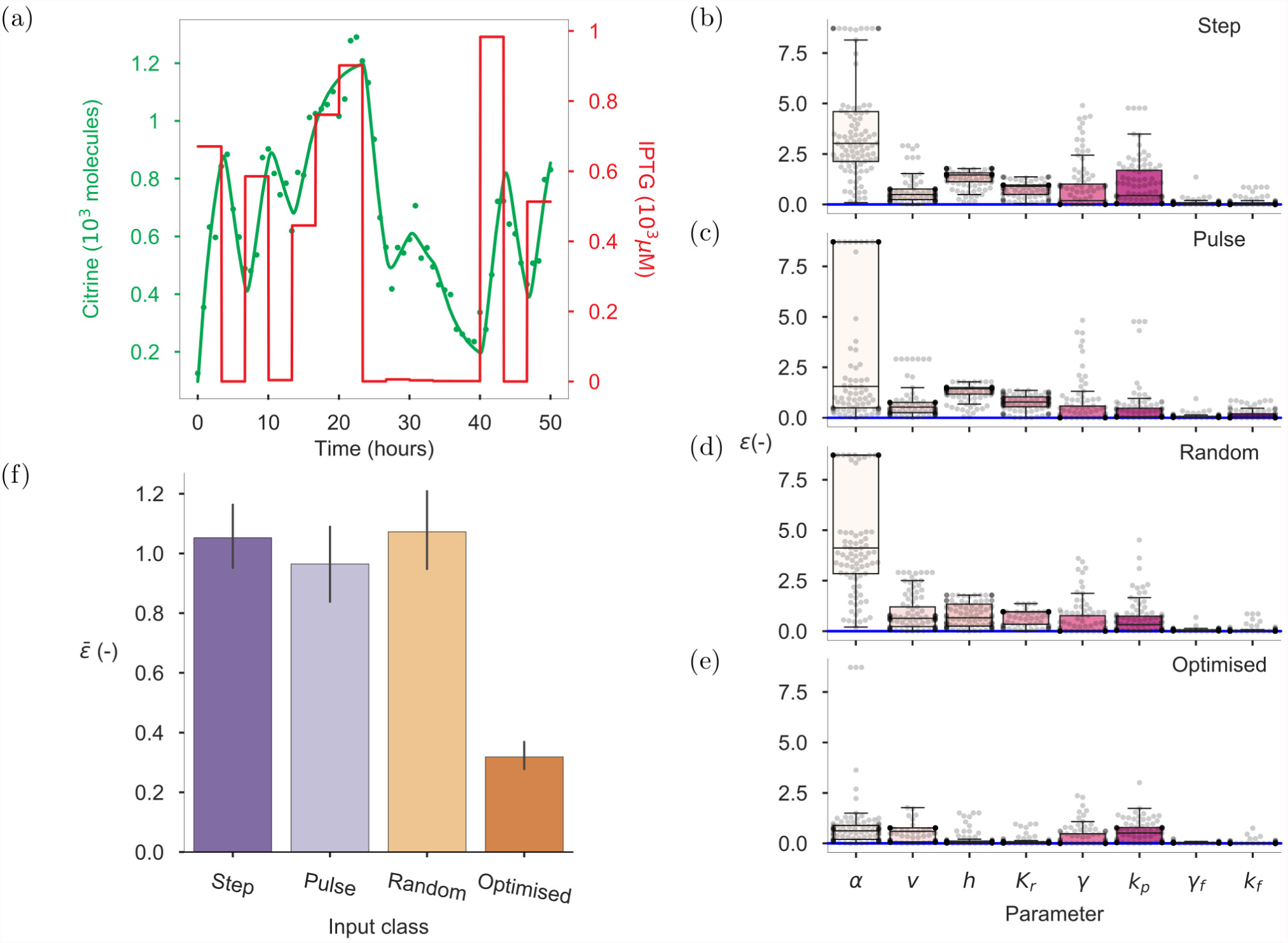
Comparison of the informative content of different input classes for model identification. (a) Example of an optimally designed input (red line) applied to M_IP,r_ to simulate Citrine dynamics (green circles). The output of the system, upon calibration to the Citrine dynamic data, is shown as a green solid line. Box plots, overlaid with swarmplots, of the relative error (ε) of parameter estimates for step (b), pulse (c), random (d) and optimised (e) inputs. (f) Bar plot of the average relative error 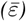 provided by each input class.

### D. Optimal Input Design (OID) enhances model calibration

We next test the improvement in the accuracy of parameter inference enabled by optimally designed experimental schemes. Having fixed the duration of the experiment, the sampling frequency and the switching times in the stepwise optimised input (Methods, section IV-D), we cast OID as a constrained optimisation problem that searches for the IPTG concentrations, (i.e. steps amplitude) that maximise the experimental information. We quantify information as the determinant of the Fisher Information Matrix 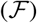 [3], [8]. This corresponds to the adoption of the highly popular D-optimality criterion [9]. To compare with the intuition-driven classes of input, we design *N_j_* = 100 optimised stimulation profiles (see Fig. 4a for an example), apply them to *M_IP,r_* to obtain pseudo-data and solve the parameter estimation problem. The results show that the use of optimised inputs leads to a marked reduction in ε when compared to experience-based stimulation patterns (Fig. 4a-e). The improvement in the accuracy of parameter estimates, noticeable for the poorly identifiable parameters *α*, *v* and *γ*, translates in a 69% reduction in 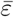 for the optimally designed input over the intuition-driven counterparts.

## III. DISCUSSIONS

Streamlining the inference of predictive mathematical models would foster their systematic use in Synthetic Biology. Here, by comparing the informative content of different input classes, we highlight optimal experimental design as a key strategy towards accurate and efficient model calibration. This conclusion was drawn considering the calibration of a deterministic model for the orthogonal, inducible promoter designed by Gnügge *et al.* [7]. We choose to focus on an inducible promoter for the key role these parts play in Synthetic Biology. Furthermore, it is commonly believed that the low complexity of synthetic promoters helps the experimentalist with the definition of informative experimental schemes based on intuition only. Our analysis clearly shows that this is not the case (Fig. 4f).

To compare the informativeness of the different input classes, we first retrieve the model structure (*M_PLac_*) proposed by the authors in [7]. The observed gap between the numerical and empirical transition-region of the dose-response curve encouraged us to attempt a model refinement by re-calibrating *M_PLac_*. We frame model inference as a multi-experimental fitting problem and, unlike Gnügge and colleagues, address it using cross validation. While *M_PLac,r_* yields a 56% improvement in the fitting over *M_PLac_* (Fig. 2c), only speculations can be made on the cause of this difference. Coherent with the *tenet* that the informative content of stimulation patterns depends on the (*a priori* unknown) dynamic properties of the system under investigation [5], we included step, pulse, ramp and random inputs in the pseudo-experiments. The match between model predictions from *M_IP,r_* and pseudo-experimental data from *M_PLac,r_* suggests that the dynamics of the latter can be fully recapitulated by the former (Fig. 3). This further supports using *M_IP,r_* as a representative model of the true biological system when comparing the informativeness of different classes of inputs for model calibration.

We find that experiments with optimised inputs provide more accurate parameter estimates than intuition-driven inputs (Fig. 4f). However, it is important to note that the lower average error provided by the optimised input does not imply that all parameter estimates improve. This is evident in our results; for example, pulse inputs allow attaining a narrower *ε* distribution for *k_p_* (Fig. 4b-d). Nevertheless, optimally designed inputs help tackling practically identifiability issues affecting some of the parameters (Fig. 4b-d).

We remark that the *a posteriori* analysis of the convergence curves of the input optimisation (results not shown) suggests that the *ε* we report should be considered an upper bound for the attainable improvement due to OED, rather than a precise estimate.

Taken together, these results suggest that a combination of *in silico* and experimental tools has the potential to significantly improve our ability to identify reliable and predictive models of biological systems and eventually enable the development of a Model-Based Biosystems Engineering framework in Synthetic Biology.

## IV. METHODS

### A. Generating Pseudo Experimental Data for the identification of M_I P,r_

To re-calibrate parameter values in *M_I P_*, and obtain *M_IP,r_*, we choose to simulate the response of *M_PLac,r_* to step, pulse, ramp and random inputs over 3000-minute long experiments. For each of these 4 input classes we define a generating function; we then design 3 inputs for each class. Step inputs are obtained using:

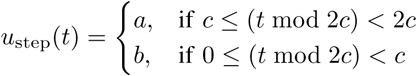

where *a*, *b* and *c* are set to [5 *µ*M, 0 *µ*M, 250 min] respectively for the first of the three time-profiles (Fig. 3A), [10 *µ*M, 0 *µ*M, 500 min] for the second and [1000 *µ*M,10 *µ*M, 500 min] for the third.

To obtain pulse inputs we use the following definition:

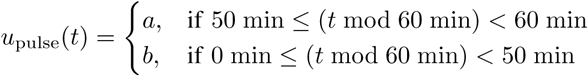

where *a*, *b* are set to [10 *µ*M, 5 *µ*M] for the first time-profile, [100 *µ*M,10 *µ*M] for the second input (Fig. 3B) and [1000 *µ*M, 600 *µ*M] for the third.

As generating function of the ramp input we use:

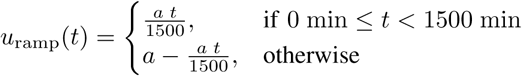

where *α* is set to 10 *µ*M, 100 *µ*M (Fig. 3C) and 1000 *µ*M for each of the three inputs generated for this class. It should also be noted that a Zero Order Holder filter with a window of 60, 150 and 250 min was applied to the first, second and third input respectively.

Finally, the pseudo-random inputs are defined as:

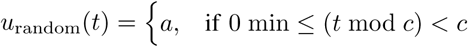

where *a*, *c* are set to [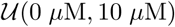, 60 min] for the first time-profile (Fig. 3D), [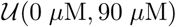,150 min] for the second and [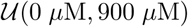, 250 min] for the third.

In all simulations, we add a 5% Gaussian noise and assign the initial conditions of the system to the steady state values derived from a 24 hour simulation of *M_PLac,r_* with 0 *µ*M IPTG as the input. All experiments are simulated in `AMTGO2` [6] and Citrine is sampled every 5 minutes. For more details on these procedures we refer the reader to our GitHub repository [10].

### B. Generating Pseudo Experimental Data for the comparison of input classes

The inputs we used to compare the informative content of different stimuli were defined as follows:

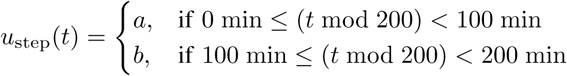

where, for each of the *N_j_* inputs, *a* and *b* are two random values extracted from 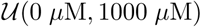.

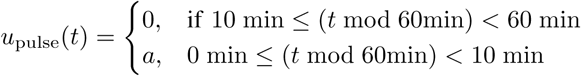

where *a* is drawn from 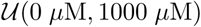.

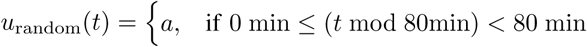

where *a* is drawn from 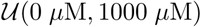

In all simulations, we add a 5% Gaussian noise and set the initial conditions of the system to the analytical steady-state of *M_IP,r_* with IPTG equal to 0 *µ*M; all experiments are simulated in `AMTGO2` [6] and Citrine is sampled every 5 minutes. For more details on these procedures we refer the reader to our GitHub repository here [10].

### C. Parameter Estimation

Parameter estimation was formulated as a non-linear optimisation problem, whose objective is to identify the parameter values that minimise a scalar measure of the distance between model predictions and (pseudo) experimental data. We use the weighted least squares as a cost function, with weights set to the inverse of the experimental noise. To solve the optimisation problem, we rely on eSS [11]: a hybrid method that combines a global and a local search to speed up convergence to optimal solutions. In the initial phase, eSS explores the space of solutions, then, as local search, the algorithm employs the nonlinear least squares solver. To strengthen the predictive capabilities of the calibrated models, we use cross validation in the identification of *M_PLac,r_* and *M_IP,r_*. In both cases, the available experimental datasets are randomised and split into training (66%) and test (33%) sets. Parameter estimation is run on the training set starting from 100 initial guesses for the parameter vector. The latter are obtained as latin hypercube samples within the allowed boundaries for the parameters. Among the optimal solutions, the one that minimises the SSE on the test set is selected as the vector of parameter estimates. It is worth noting that, when comparing the informative content of different input classes, parameter estimation was not performed using cross validation. Details on the allowed bounds for the parameters and the scripts used for parameter estimation are provided in the GitHub repository [10].

### D. Optimal Experimental Design

To reflect wet-lab experimental constraints, we fix the sampling times (1 every 5 minutes) and the experiment duration (3000 minutes). We further set the initial condition to the steady-state in absence of induction. As a result, we restrict the optimisation to identifying the input (IPTG) time profile that maximises the information yield of the experiment. Here, information is quantified as a metric defined on the Fisher Information Matrix 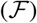 [3], [8]:

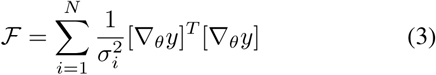

where *y* is the observable (Citrine) and 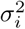 represents the variance of the signal at the *i*^th^ sampling instant.

The 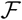 sets a lower bound on the variance of the parameter estimates through the Cramér-Rao inequality:

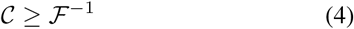

where 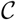 is the covariance matrix. Intuitively, as the eigen-values of the 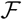 are related to the inverse of parametric variances, attempting to maximise the determinant of 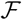 (D-optimality) corresponds to minimising the product of the parametric variances.

In order to find the most informative input (*u*\*), we formulate MBOED as an optimal control problem and search for:

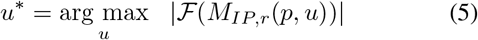

where *p* is the parameter vector. We use Differential Evolution (*DE*) [12], a global optimisation method featuring good convergence properties and suitable for parallelisation, to solve the optimisation problem. We empirically [13] set the population size, crossover threshold and differential weight to 150, 0.3 and 0.5, respectively and adopt the strategy randto-best/1/exp.

## V. CONCLUSIONS AND FUTURE WORK

In this study we highlight MBOED as a key strategy for the accurate calibration of mathematical models of biological parts in Synthetic Biology. Our *in-silico* results suggest that optimally designed input profiles substantially improve the predictive ability of the inferred models, outperforming intuition-driven stimuli. While further studies are needed to explore the scalability of the computational cost for systems of higher complexity, we propose that combining flexible experimental platforms (e.g. microfludics) and MBOED will enable the widespread adoption of mathematical models in Synthetic Biology. Beyond the required *in vivo* validation, our results encourage efforts towards the implementation of platforms to automate model calibration, in which MBOED and *in vivo* experiments are combined in an identification loop.

## ACKNOWLEDGMENTS

We would like to thank Prof. Jürge Stelling and Ms. Dharmarajan Lekshimi for clarifications on the analysis of experimental data and on the model implementation in [7]. We thank Mr. Alastair Hume for his contribution to the simulation code used in this study. We are further grateful to Prof. Diego di Bernardo and his lab for insightful discussions.

